# Predictability of Genetic Interactions from Functional Gene Modules

**DOI:** 10.1101/049627

**Authors:** Jonathan H. Young, Edward M. Marcotte

## Abstract

Characterizing genetic interactions is crucial to understanding cellular and organismal response to gene-level perturbations. Such knowledge can inform the selection of candidate disease therapy targets. Yet experimentally determining whether genes interact is technically non-trivial and time-consuming. High-fidelity prediction of different classes of genetic interactions in multiple organisms would substantially alleviate this experimental burden. Under the hypothesis that functionally-related genes tend to share common genetic interaction partners, we evaluate a computational approach to predict genetic interactions in Homo sapiens, Drosophila melanogaster, and Saccharomyces cerevisiae. By leveraging knowledge of functional relationships between genes, we cross-validate predictions on known genetic interactions and observe high-predictive power of multiple classes of genetic interactions in all three organisms. Additionally, our method suggests high-confidence candidate interaction pairs that can be directly experimentally tested. A web application is provided for users to query genes for predicted novel genetic interaction partners. Finally, by subsampling the known yeast genetic interaction network, we found that novel genetic interactions are predictable even when knowledge of currently known interactions is minimal.

## Introduction

Determining the genetic interactions in an organism provides a basis for understanding how the role of a gene is influenced by the action of any other gene. By definition, two or more genes interact when combining variants of each gene produces a significantly pronounced phenotype when compared to the phenotypes of individual variants [Mani et al., 2008, Baryshnikova et al., 2013]. The applications of exploiting such interactions extend to drug target discovery. Strategies such as targeting genes that interact with cancer-specific mutations have been proposed and reviewed extensively [Ashworth et al., 2011, Fece de la Cruz et al., 2015] and have led to clinical trials [Fong et al., 2009]. Because experimental determination of genetic interactions involves examining all possible pairs from a group of genes, practical difficulties arise when a comprehensive interaction map of an entire organism is desired. Multicellular organisms present the challenge of various differentiated cell types, each having potentially differing genetic interactions. Moreover, there are different kinds of genetic interactions, ranging from those based on growth effects to other phenotypic effects. There exists a need to either reduce the search space for testing genetic interactions or to reliably predict them. Here, we evaluate a computational approach to predict and validate different types genetic interactions across multiple organisms.

Previous studies to predict genetic interactions leveraged existing sources of biological information. Integration of biological features in yeast (i.e. gene co-expression, protein interaction and function) and their associated network topological properties guided the training of probabilistic decision trees to predict synthetic sick or lethal (SSL) interactions [Wong et al., 2004]. In a similar vein, an ensemble classifier was trained on a set of 152 genetic interaction-independent features to predict SSL in yeast [Pandey et al., 2010]. Compiling multiple biological features has also been extended to more than one organism. By considering the orthologous gene pairs among yeast, fly and worm, features such as functional annotation were used to train a logistic regression model to predict a genome-wide map of genetic interactions [Zhong and Sternberg, 2006]. Alternatively, studies have also explored network-based approaches for genetic interaction prediction. Novel SSL interactions were predicted by way of a diffusion kernel on a network of known SSL gene pairs [Qi et al., 2008]. Interrogating functional gene networks that were constructed from integration of biological data from literature have proven useful in predicting modifier genes in yeast and worm [Lee et al., 2010]. Many of these approaches have focused on a single genetic interaction type in a single organism.

Here, we examine an algorithm to predict multiple types of genetic interactions across diverse organisms based on the hypothesis that genes strongly participating in shared functions also share common genetic interaction partners. Our approach relies on a functional gene network for a given organism and knowledge of known genetic interactions of a particular type. We tested our approach on three organisms - human (*Homo sapiens*), fly (*Drosophila melanogaster*), and yeast (*Saccharomyces cerevisiae*) - and found predictability across different types of genetic interactions. We also investigated how some interactions are enriched in yeast and human gene modules, specifically protein complexes, and the degree to which genetic interactions need to experimentally determined before enrichment can be found.

## Materials and Methods

For various classes of genetic interactions in human, fly, and yeast, a list of genes and each of their known genetic interaction partners were assembled. A gene and its known interaction partners are collectively referred to as a “seed set.” Receiver operating characteristic (ROC) analysis was performed to quantify whether the interaction partners of any given gene are clustered in the organism’s functional gene network. Specifically, for every group of interaction partners of a gene, a score vector consists of entries that are sums of functional network edge weights between each gene in the network to the interaction partners. Because there are no self-edges in the network, leave-one-out cross-validation is carried out on the known interaction partners. An accompanying label vector indicates whether each gene in the network is indeed an interaction partner. The two vectors yield a ROC curve and the corresponding area under the curve (AUC). A seed set’s AUC is the measure of how tightly connected the interaction partners are in the functional network and therefore how predictive the seed set is for novel interactions [Lee et al., 2010]. None of the known genetic interactions used for prediction were contained in the functional gene network.

Enrichment of genetic interactions within yeast and human protein complexes was calculated with a binomial model defined as 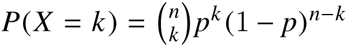, where the background probability *p* equals the proportion of all possible gene pairs that are genetically interacting. The number of trials *n* is the number of possible gene pairs in the complex, and *k* equals the number of interacting pairs in the protein complex.

### Statistical Analysis

If *k* is the number of genetic interactions within a protein complex, then the corresponding *p*-value is *P*(*X* ≥ *k*) according to a binomial model as previously described, with control of FDR at 5% through the Benjamini-Hochberg procedure [Benjamini and Hochberg, 1995]. Seed sets with AUC ≥ 0.9 were considered highly predictive of novel genetic interactions.

### Data Availability

All genetic interactions were downloaded from version 3.4.130 of BIOGRID [Stark et al., 2006]. Organism-specific functional gene networks were downloaded for human [Lee et al., 2011], fly [Shin et al., 2015], and yeast [Lee et al., 2007]. Previous studies served as sources of protein complexes for yeast and human [Hart et al., 2007, Ruepp et al., 2010]. Python code using the Matplotlib [Hunter et al., 2007], scikit-learn [Pedregosa et al., 2011], and *mygene* [Wu et al., 2012] libraries is available at https://bitbucket.org/youngjh/genetic_interact. All network visualizations were produced in Cytoscape [Shannon et al., 2003]. A supplementary web page at http://marcottelab.org/Genetic_Interact/ allows users to query a gene of interest. If the gene has known genetic interaction partners that are predictive, then the functional network cluster is displayed. Raw data files listing the seed sets with AUC ≥ 0.7 are also available.

## Results

We sought to determine whether clusters of functionally related genes, for example genes *A-E* in Figure 1, are predictive of genetic interactions. In this example, genes *A* and *C-E* are known to share genetic interactions with gene *X*, and our hypothesis would suggest gene *B* as a novel interaction partner of *X*. Our method identifies predictive clusters by leave-one-out cross-validation and receiver operating characteristic (ROC) analysis; when applied to the network in Figure 1, each of genes *A* and *C-E* are individually withheld as known interaction partners one at a time and predicted back with high recall. Subsequently, gene *B* is a novel high-confidence predicted interaction partner of *X*. The approach described here was evaluated for several classes of phenotypic and growth-based genetic interactions in human, fly and yeast.

**Figure 1:**
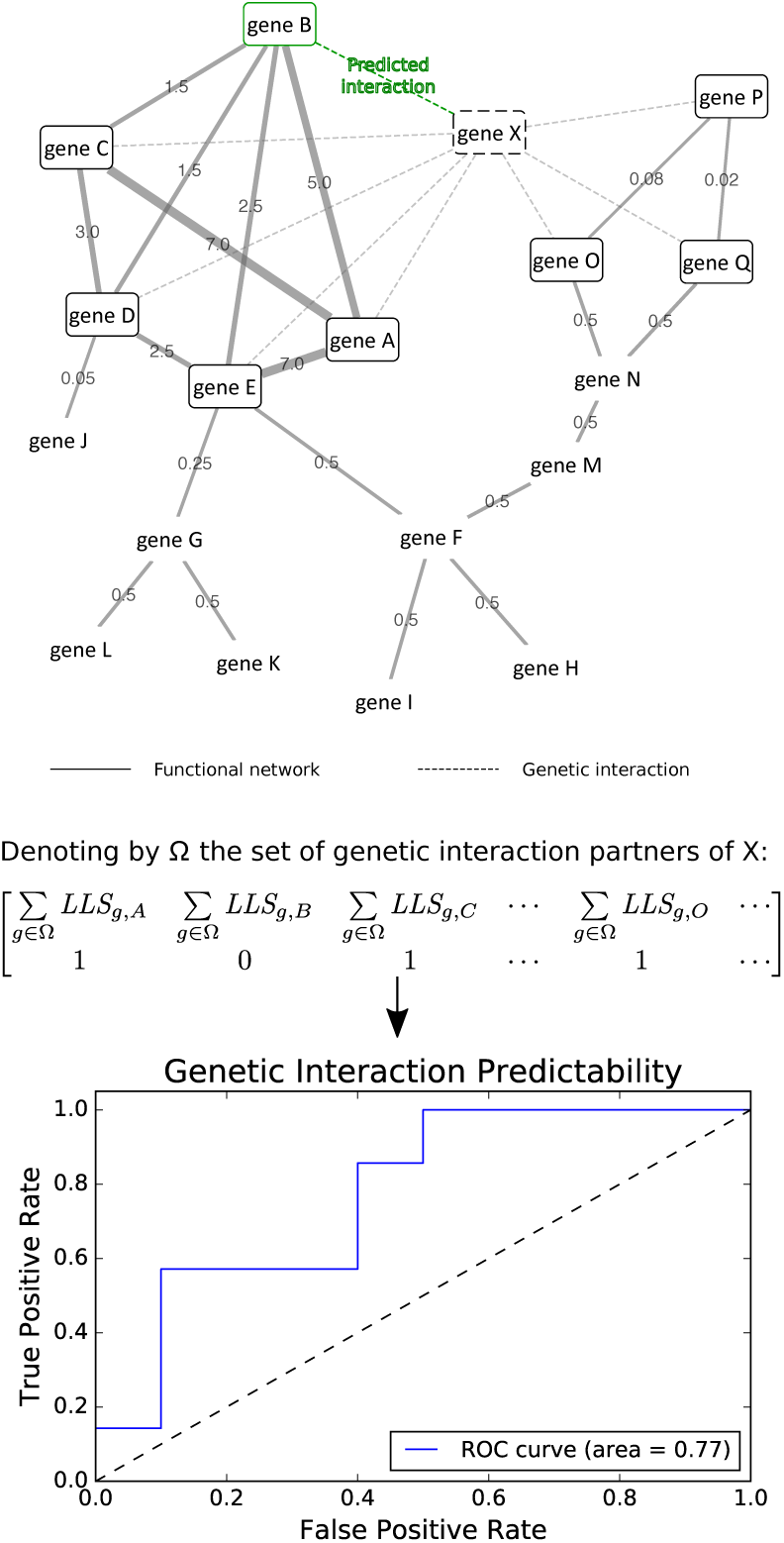
Genetic Interaction Prediction. Dashed edges indicate known genetic interactions. Solid edges connect genes that participate in the same biological process, with log-likelihood (LLS) scores as edge weights reflecting the degree of confidence in the genes’ shared functionality. Genes *A, C-E* are genetic interaction partners of gene *X* and members of a functional net cluster; then the remaining cluster member, gene *B*, is a predicted interaction partner of gene *X* as well. Candidate clusters are evaluated by first assigning scores to each gene in the network by summing the edge weights, as shown in the first row of the matrix. *LLS*_*g,A*_ denotes the log-likelihood score between genes *g* and *A*. The second row is populated with binary labels indicating whether the gene is a known interaction partner of *X*. In this fashion, a ROC curve is constructed to yield an AUC.

### The human functional gene network is predictive for phenotype-based genetic interactions

As shown in Figure 2A, our method demonstrated high performance in predicting phenotypic enhancing and suppressing human gene pairs. In these interactions, a double mutant has an enhanced or suppressed phenotype (other than growth) in comparison to either of the single mutants. The plots for phenotypic enhancement and suppression in Figure 2A display the performance of seed sets, each of which are defined as a group of known phenotypic enhancing or suppressing partners of a particular gene. There are 238 phenotypic enhancement seed sets, of which 30 have AUC ≥ 0.9. Similarly, 36 of 215 phenotypic suppression seed sets have AUC ≥ 0.9. The AUC is the area under the receiver operating characteristic (ROC) curve that measures how well the known interaction partners rank in our leave-one-out cross-validation scheme. Those that are not predictive are the ones with AUC = 0.5, indicating that their predictability is no better than random. For the most part, seed sets are either at least moderately predictive, or not at all.

**Figure 2:**
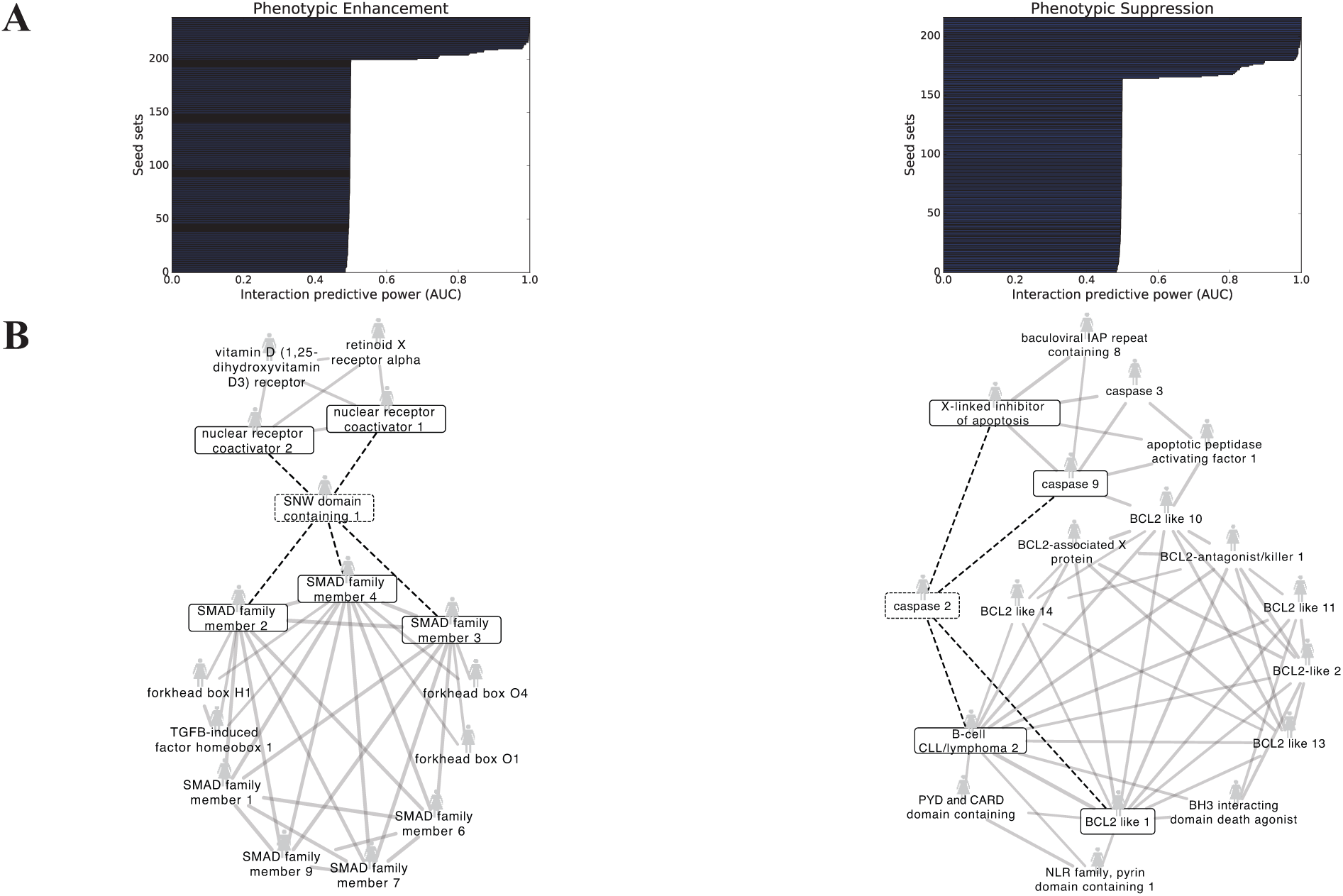
Predictive functional net clusters yield novel phenotypic enhancing and suppressing human gene pairs. (A) Each horizontal bar represents the set of known genetic interaction partners of a specific human gene; each of these sets is referred to as a “seed set.” High AUC scores indicate that the interaction partners participate together in a cluster in HumanNet, the human functional gene network. Therefore, other members of the cluster are predicted as novel interaction partners. (B) Shown are two examples of well-defined HumanNet clusters that are highly predictive for phenotypic enhancement (left) and suppression (right), with the known interactions from the seed set denoted by the boxed genes and dashed edges.

Shown in Figure 2B are illustrative seed sets with high predictability that form well-defined clusters in the human functional gene network, HumanNet. For clarity, only functional network edges with log-likelihood scores (LLS) above 3.0 are shown. Furthermore, HumanNet genes are shown only if they connect to at least 2 of the known genetic interaction partners. The seed set consisting of the SNW domain containing 1 in phenotypic enhancement with members of the SMAD family and nuclear receptor coactivators yielded an AUC of 0.91. The prediction is that the SNW domain containing 1 also phenotypically enhances with other members of the SMAD family along with members of the forkhead box. In the phenotypic suppression case, we find that known phenotypic suppressors of caspase 2 are tightly functionally linked with members of the BCL2-like family, among other genes. With a resulting AUC of 0.90, these BCL2-like genes are expected to participate in phenotypic suppression with caspase 2.

### Fly phenotypic enhancement and suppression interactions are predicted from functional net clusters

Similar to the human case, the fly functional network FlyNet is particularly predictive of phenotypic enhancement and suppression, as shown in Figure 3. A larger proportion of the seed sets are predictive than in the human case. For phenotypic enhancement, 322 out of 754 seed sets had AUC ≥ 0.9, and 398 phenotypic suppression seed sets (out of 818) met the same threshold. Figure 3B shows a well-defined gene cluster (AUC = 0.94) containing phenotypic enhancement interaction partners of seven up. From this cluster, genes involved in the sevenless signaling and the Drosophila epidermal growth factor receptor signal transduction pathways achieved high recall, and neighbor genes also involved in the same signaling pathways are expected to phenotypically enhance seven up. Turning to phenotypic suppression, several Enhancer of split genes are tightly clustered (AUC = 0.98) with known phenotypic suppressors of hairy that include the *achaete-scute* complex, thereby implicating them as additional, novel phenotypic suppressors of hairy.

**Figure 3:**
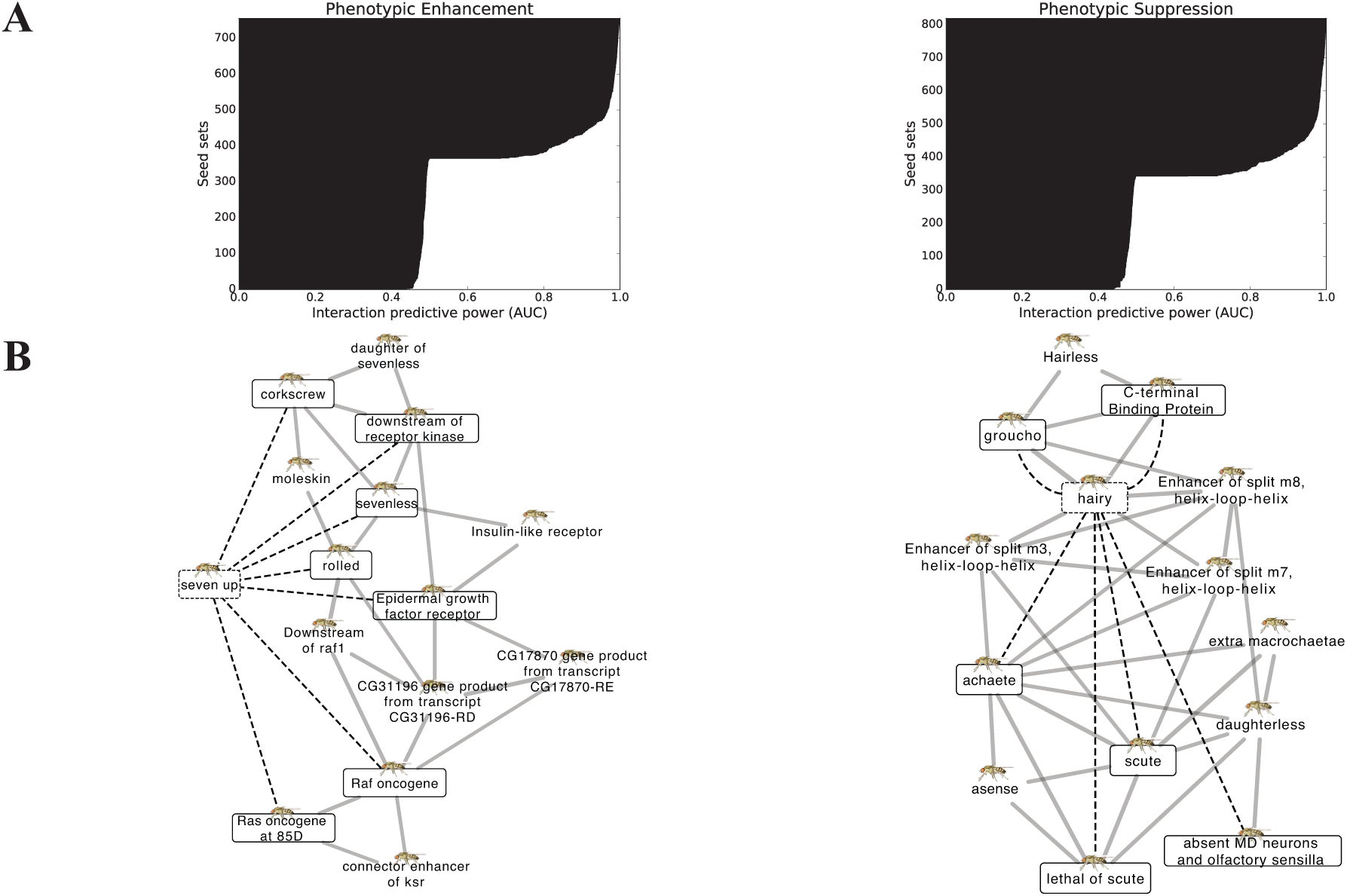
FlyNet predictability for phenotypic enhancing and suppressing genetic interactions. (A) Each horizontal bar represents a single fly gene that is known to interact with a number of other genes. (B) Predictive seed gene sets are shown for phenotypic enhancement (left) and suppression (right).

### High-confidence predictability is found in human, fly and yeast

The full range of various genetic interaction classes that were analyzed from BIOGRID are listed in Table 1. Genetic interactions were generally based on phenotypic effects or growth and lethality measurements. Each entry in Table 1 lists the number of predictive seed sets having AUC ≥ 0.9 of out the total examined. In human, our method performed well primarily for phenotypic enhancement and suppression as described above, but did not offer predictability for the dosage lethality and synthetic growth defect and rescue interactions determined to date. For fly, most of the known interactions fall into the phenotypic enhancement and suppression categories, for which high predictability was observed. Although a moderate number of fly dosage rescue interactions are known, no predictive seed sets were found. In both human and fly, several classes of interactions have not been extensively determined and thus were untested in our prediction scheme.

**Table 1:** **Predictive power of functional networks across different genetic interactions**.

|  | *H. sapiens* | *D. melanogaster* | *S. cerevisiae* |
| --- | --- | --- | --- |
| Dosage Growth Defect | Not tested | Not tested | 176 / 1146 |
| Dosage Lethality | 2 / 108 | Not tested | 116 / 689 |
| Dosage Rescue | 5 / 65 | 0 / 144 | 203 / 1358 |
| Phenotypic Enhancement | 30 / 238 | 322 / 754 | 287 / 1958 |
| Phenotypic Suppression | 36 / 215 | 398 / 818 | 223 / 1751 |
| Synthetic Growth Defect | 4 / 445 | 1 / 5 | 576 / 3417 |
| Synthetic Rescue | 2 / 131 | 5 / 26 | 218 / 2089 |
| Synthetic Lethality | Not tested | Not tested | 221 / 2706 |
| Negative Genetic | Not tested | Not tested | 65 / 4618 |
| Positive Genetic | Not tested | Not tested | 55 / 3586 |

For each fraction, the numerator indicates the number of seed sets with AUC ≥ 0.9 and the denominator equals the total number of seed sets tested.

Our method also performed well in most of the interaction categories for *S. cerevisiae* (Table 1, Supplementary Figure S1). Notably, negative and positive genetic interactions fared poorly as few predictive seed sets were identified, even though most of the experimentally determined interactions in yeast fall into these categories.

### Protein complexes inform trends of genetic interaction predictability

With genetic interactions predicted across multiple organisms, it was natural to investigate their evolutionary conservation. In particular, if a protein complex were enriched in genetic interactions, then perhaps a homologous protein complex would also exhibit similar enrichment. We found enrichment of various types of interactions within yeast protein complexes, but none thus far for human. Therefore, instead the problem shifted to identifying the degree to which genetic interactions must be determined in order to find enrichment, and therefore predictability. Using yeast as a test case, simulations successively withheld increasing proportions of genetic interactions, with enrichment within yeast protein complexes computed at each point. The interaction types considered were negative and positive genetic, and synthetic growth defect and lethality. As shown in Figure 4, when withholding genes with a genetic interaction degree (the number of interacting partners of a certain gene) of more than 5, corresponding to withholding >90% of synthetic growth defect and >80% of synthetic lethality pairs, then an immediate drop-off in enrichment resulted. No such behavior was observed for negative and positive genetic interactions, for which enrichment linearly decreased as a function of the withheld proportion. Similarly, when removing interacting pairs at random, there was a steady decrease in the number of significantly enriched complexes among all types. Finally, when withholding pairs under a degree cutoff, there was also no point beyond which enrichment failed to be found (Supplementary Figure S2).

**Figure 4:**
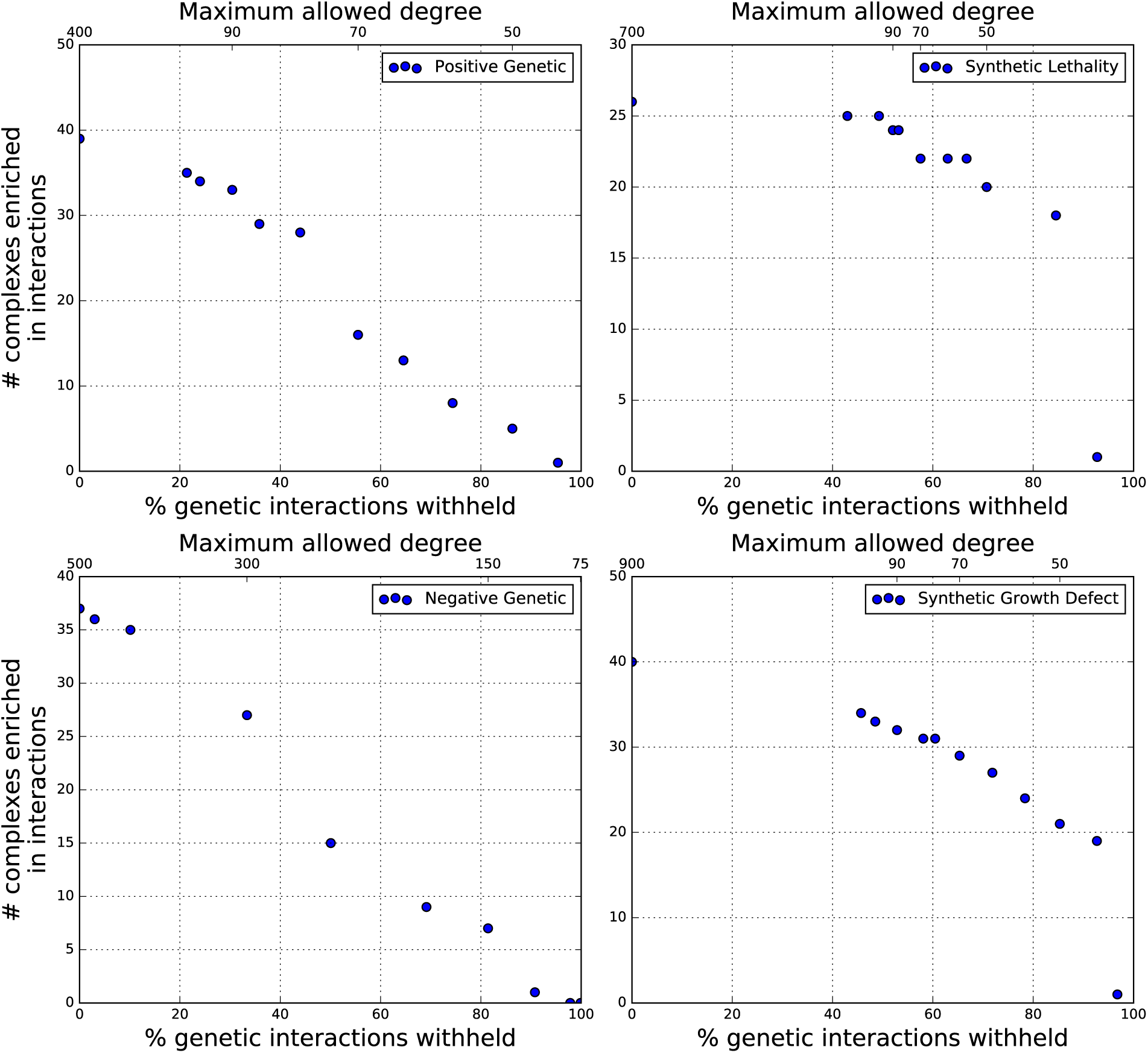
Predictability of genetic interactions can be found even when known interactions are sparse. By successively withholding known yeast genetic interactions according to each gene’s interaction degree (e.g. number of interaction partners), enrichment and therefore predictability is still detectable when information of known interactions is minimal. This effect is especially pronounced for synthetic growth defect and lethality, provided genes possess sufficiently high interaction degree.

## Discussion

Our results demonstrate that various classes of genetic interactions in different organisms can be successfully predicted based on the hypothesis that functional gene clusters tend to share genetic interaction partners. For *S. cerevisiae* in particular, predictability was obtained whether the genetic interaction type was based on growth effects or non-growth phenotype-based measurements (i.e. phenotypic suppression). Interestingly, our method did not yield predictability for negative and positive genetic interactions, which happen to be the interaction types for which most of the pairs have been tested [Costanzo et al., 2010]. While the range of predictable genetic interaction classes for human and fly were limited to phenotypic enhancement and suppression, we believe that this is probably due to the sparsity of known genetic interactions for these organisms. In this study, the source of known genetic interactions, BIOGRID, had over 150000 yeast gene pairs but only ~2800 pairs for fly and ~1500 for human. As shown in Table 1, many types of genetic interactions could simply not be tested for fly and human.

This sparseness of experimentally-determined genetic interactions, especially in human, led to the lack of enrichment in gene modules such as protein complexes. In our simulations of withholding genetic interacting pairs, we expected that regardless of the interaction type, there would be a point after which no enrichment would be found. Thus, it was surprising that negative and positive genetic interactions exhibited a linear decrease in enrichment, regardless of how the pairs were withheld (by degree or at random). On the other hand, the enrichment signal in synthetic growth defect and lethality is sensitive to the interaction degree, as there was a steep drop-off when most of the interaction pairs were withheld. In the negative and positive genetic networks, there appears to sufficient genetic interaction density such that even when high numbers of interacting pairs are withheld, enrichment under a binomial model can still be found. By extrapolating to the human case, a modest increase in the number of screened human gene pairs is likely to dramatically increase the ability to predict additional genetic interactions, especially for synthetic growth defect and lethality where the genes have multiple interaction partners.

Similar to previous genetic interaction prediction approaches [Qi et al., 2008, Zhong and Sternberg, 2006], our algorithm requires knowledge of known experimentally determined genetic interactions. While other studies proceed without such requirements, the assimilation of a host of biologically annotated features are still necessary for their prediction method [Pandey et al., 2010, Wong et al., 2004]. In contrast to the aforementioned studies, our methodology systematically examined more than one class of genetic interaction and was successfully applied to multiple eukaryotic organisms, thereby generalizing results from a previous study by Lee et al. [Lee et al., 2010]. Since the detection of tightly connected sets of nodes in a network is central to our method, further avenues for exploration perhaps include investigating methods such as graph clustering [Enright et al., 2002] or community detection algorithms [Fortunato, 2010], though these algorithms lack built-in validation. It would also be interesting to explore using tissue-specific gene networks instead of a single integrated functional gene network for more targeted predictions [Greene et al., 2015].

As one major goal of any genetic interaction prediction is to at least narrow down the search space for experimentally testing genetically interacting pairs, our predictions are specifically testable experimentally, perhaps through CRISPR-Cas9 for human cells [Wong et al., 2016]. We also contribute to available prediction methodologies for suggesting genetic interactions as candidate therapeutic targets. Ultimately, we demonstrate the power of leveraging knowledge of known genetic interactions and integrated biological information in functional gene networks to predict novel genetic interactions from single-cell to multicellular organisms.

## Acknowledgments

E.M.M. acknowledges funding from the National Institutes of Health, the National Science Foundation, the Cancer Prevention and Research Institute of Texas, and the Welch Foundation (F1515). The authors thank Kevin Drew for assistance with web server setup.

## Conflict of interest

none declared.

